# A unified open-source platform for multimodal neural recording and perturbation during naturalistic behavior

**DOI:** 10.1101/2023.08.30.554672

**Authors:** Jonathan P. Newman, Jie Zhang, Aarón Cuevas-López, Nicholas J. Miller, Takato Honda, Marie-Sophie H. van der Goes, Alexandra H. Leighton, Filipe Carvalho, Gonçalo Lopes, Anna Lakunina, Joshua H. Siegle, Mark T. Harnett, Matthew A. Wilson, Jakob Voigts

**Affiliations:** Department of Brain and Cognitive Sciences, MIT, Cambridge, MA, USA; Picower Institute for Learning and Memory, MIT, Cambridge, MA, USA; Open Ephys Inc. Atlanta, GA, USA; Dept. of Electrical Engineering, Polytechnic University of Valencia, Valencia, Spain; McGovern Institute for Brain Research, MIT, Cambridge, MA, USA; Open Ephys Production Site, Lisbon, Portugal; NeuroGEARS Ltd, London, UK; Allen Institute for Neural Dynamics, Seattle, Washington, USA; HHMI Janelia Research Campus, Ashburn, VA, USA

## Abstract

Behavioral neuroscience faces two conflicting demands: long-duration recordings from large neural populations and unimpeded animal behavior. To meet this challenge, we developed ONIX, an open-source data acquisition system with high data throughput (2GB/sec) and low closed-loop latencies (<1ms) that uses a novel 0.3 mm thin tether to minimize behavioral impact. Head position and rotation are tracked in 3D and used to drive active commutation without torque measurements. ONIX can acquire from combinations of passive electrodes, Neuropixels probes, head-mounted microscopes, cameras, 3D-trackers, and other data sources. We used ONIX to perform uninterrupted, long (∼7 hours) neural recordings in mice as they traversed complex 3-dimensional terrain. ONIX allowed exploration with similar mobility as non-implanted animals, in contrast to conventional tethered systems which restricted movement. By combining long recordings with full mobility, our technology will enable new progress on questions that require high-quality neural recordings during ethologically grounded behaviors.

---

There is a growing recognition that, to maximize their explanatory power, neural recordings must be conducted during normal animal behavior. From the recent discovery that motor actions can dominate the activity of brain regions that were believed to be predominately sensory^1,2^, to findings of different learning strategies between head-fixed and freely moving subjects^3^, mounting evidence indicates that free behavior transforms the function of the nervous system. These observations are leading towards a consensus that learning^3,4^, social interactions^5,6^, sensory processing ^7,8^, and cognition^9,10^ are best addressed in animals that engaged in naturalistic behavior.

In recent years, remarkable progress has been made on methods for tracking and quantifying animal behavior^11–42^. Parallel advances in recording technologies have enabled electrophysiology^43,44^, optical imaging^45–47^, and actuation of neural ensembles^48^ in mobile animals. Still, applying these technologies, which are often bulky and require tethers, to study naturalistic behavior remains a major challenge. In larger animals like rats^49^, primates^19^, or even bats^50^, wireless systems are available. However, in mice, which are the predominant animal model system in neuroscience, recordings are limited by the weight of recording devices. For example, a 6 g wireless logger can achieve only a 70 minute long recordings^51^ and the weight of its batteries limits movement beyond slow locomotion, requiring that experiments be designed around the head torque imposed by the recording device^51^. Therefore, current technologies for mice, and similar sized species, do not allow for unencumbered motion, nor for recordings during behavior that unfolds over long periods, limiting our ability to capture neural activity during ethologically relevant behaviors.

To address this need, we developed an open-source multi-instrument hardware standard and API (Open Neuro Interface, ‘ONI’, Suppl. Figs 1,2). We then used ONI to implement a recording system called ‘ONIX’ – a modular and extendable data acquisition and behavior tracking system that greatly reduces the conflict between large-scale neural recordings and their impact on mouse behavior. The system uses a thin and light micro coaxial tether (∼0.31 mm diameter, 0.37 g/m) compared to widely used options (e.g. 3 mm diameter, 6.35 g/m, Fig. 1c), that causes minimal forces on the animal’s head (Fig. 1d). The tether simultaneously powers and transmits data (150 MB/s, equivalent to 2500 channels of spike-band electrophysiology data) to and from sensors and actuators. ONIX includes modular, miniaturized headstages (Figs. 1 & 4, Suppl. Fig.3) for passive electrical recording probes, tetrode drives^44^ (via Intan RHD and RHS chips), and Neuropixels^43^ (1.0, 2.0 probes, and BGA-packaged chips). In addition to neural activity, these headstages record 6-degrees-of-freedom head pose at ∼30 Hz via on-board sensors with sub-degree and sub-millimeter resolution (90% of jitter < 0.02 mm at 2 m distance, Suppl. Fig. 8) via a consumer-grade 3D-tracking system (HTC Vive). Real-time tracking permits the system to measure the rotation of an animal and automatically untwist the tether via a small motor (Fig. 1b, Suppl. Fig. 5) without requiring torque measurements. This approach removes the behavioral impact and time limits typically associated with tethered recordings and provides experimenters with the ability to monitor neural activity and behavior for arbitrarily long sessions in complex environments (Figs. 2 & 3).

**Figure 1:**
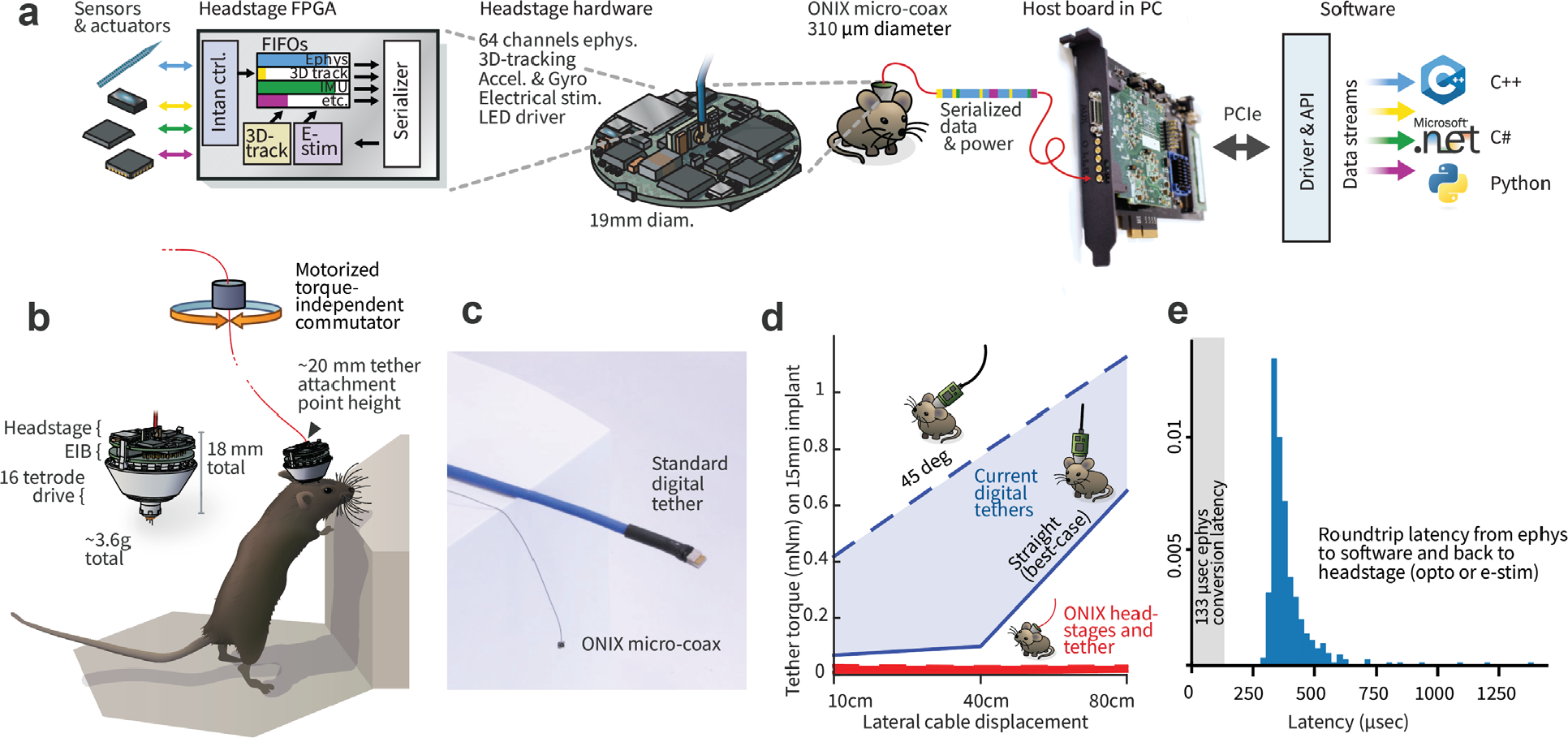
ONIX, a unified open-source platform for unencumbered freely moving recordings. **a**, Simplified block diagram of the ONI (open neuro interface), illustrated via the tetrode headstage: multiple devices all communicate with the host PC over a single micro-coax cable via a serialization protocol, making it possible to design small multi-function headstages. **b**, The integrated inertial measurement unit and 3D-tracking redundantly measure animal rotation, which drives the motorized commutator without the need to measure tether torque. Small drive implants^44^ enable low-profile implants (∼20 mm total height). **c**, The ONIX micro-coax, a 0.31 mm thin tether of up to 12 m length, compared to standard 12-wire digital tethers. **d**, Torque exerted on an animal’s head by tethers. Current tethers allow full mobility only in small arenas and in situations when the tether does not pull on the implant, while the ONIX micro-coax applies negligible torque. **e**, Performance of ONIX: With the 64 channel headstage, a 99.9% worst case closed-loop latency, from neural voltage reading, to host PC, and back to the headstage (e.g. to trigger an LED) of <1 ms can be achieved on Windows 10 (see also Suppl. Figs 6 & 7).

**Figure 2:**
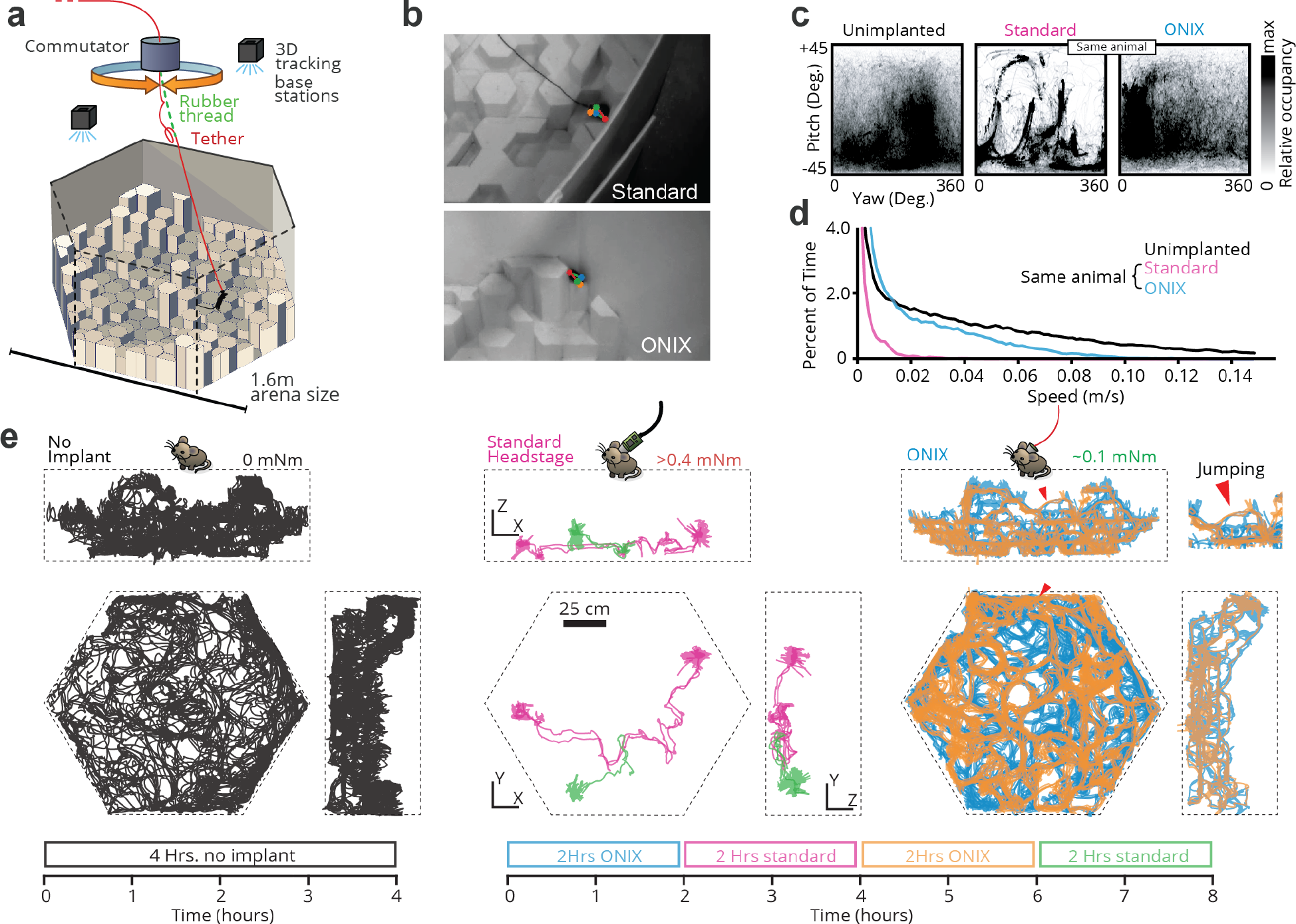
Unrestricted naturalistic locomotion behavior with ONIX. **a**, Overview of experiment: Mice were freely exploring a 3D arena made out of styrofoam pieces of varying heights. **b**, Unimplanted mice and mice with a standard tether (top) or ONIX micro coax (bottom) were tracked in 3D using multicamera markerless pose estimation^31^. **c**, Head yaw and pitch occupancies over the course of a recording. Standard tethering significantly reduces freedom of head movement relative to unimplanted mice, whereas ONIX does not. **d**, Speed distributions over the course of a recording. Standard implants significantly reduce running speed, while ONIX results in only a modest speed reduction relative to unimplanted mice. **e**, 2D projection of mouse trajectories over the course of a recording session. ONIX retains the same spontaneous exploration behavior as unimplanted mice, while standard headstages and cables greatly reduce locomotion.

**Figure 3:**
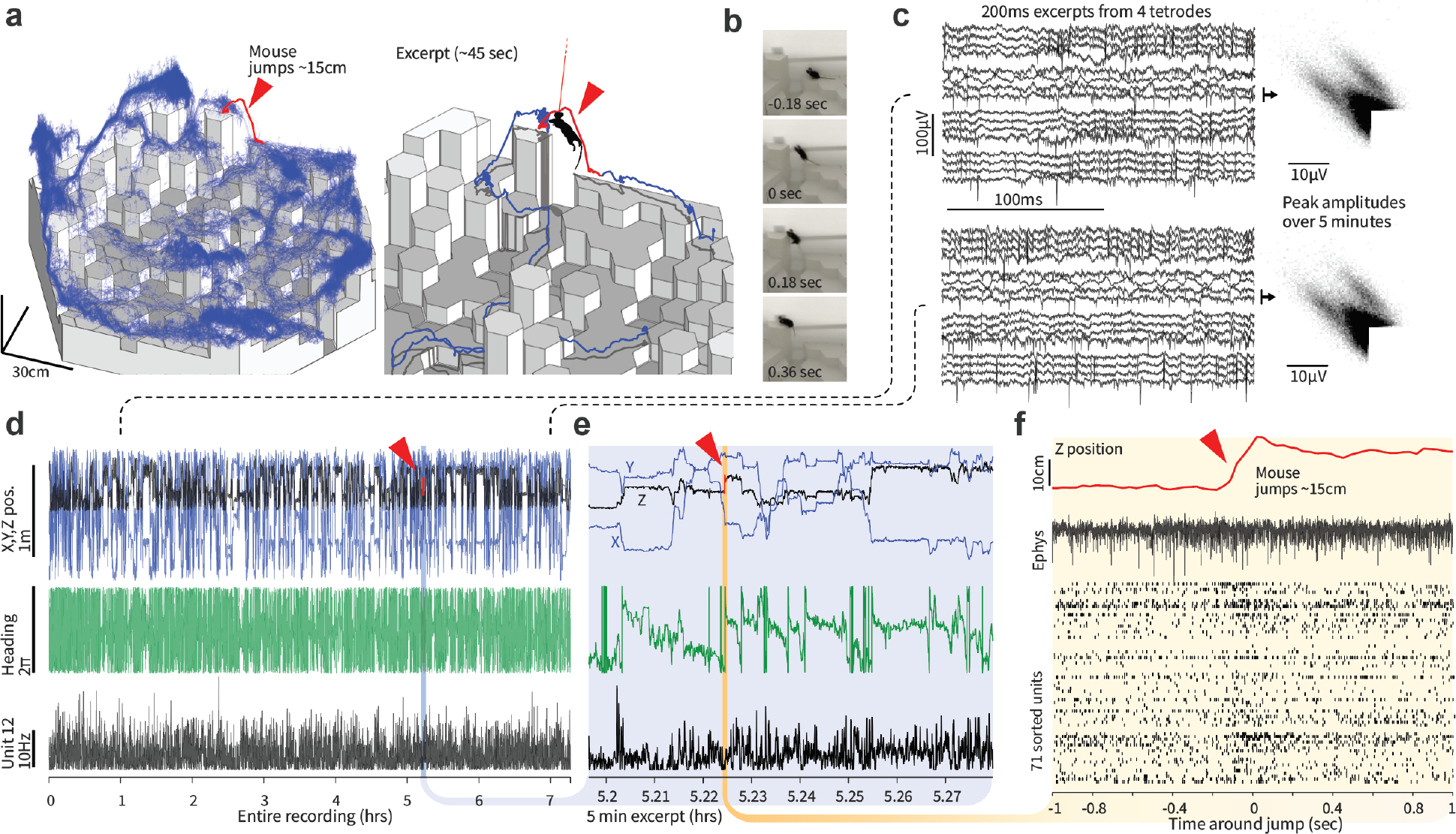
Stable long-term recordings during naturalistic locomotion. **a**, Position of one 3D-tracking sensor on the headstage during a 7.3 hour-long ONIX recording during which the mouse was free to explore the 3D arena. Red trace and excerpt show one of multiple instances of the mouse spontaneously jumping from a lower to a higher tile. **b**, Video frames of the jump (the tether is too thin to be visible at this magnification), see supplementary video 1. **c**, Raw voltages and spike peak amplitudes from the beginning (left) and end (right) of the recording. **d**, 3D-position, heading, and smoothed firing rate of entire recording. **e**, Same data as in d, for excerpt around jump. **f**, Z-position, raw voltage trace example, and sorted spikes from 71 neurons during the jump.

**Figure 4:**
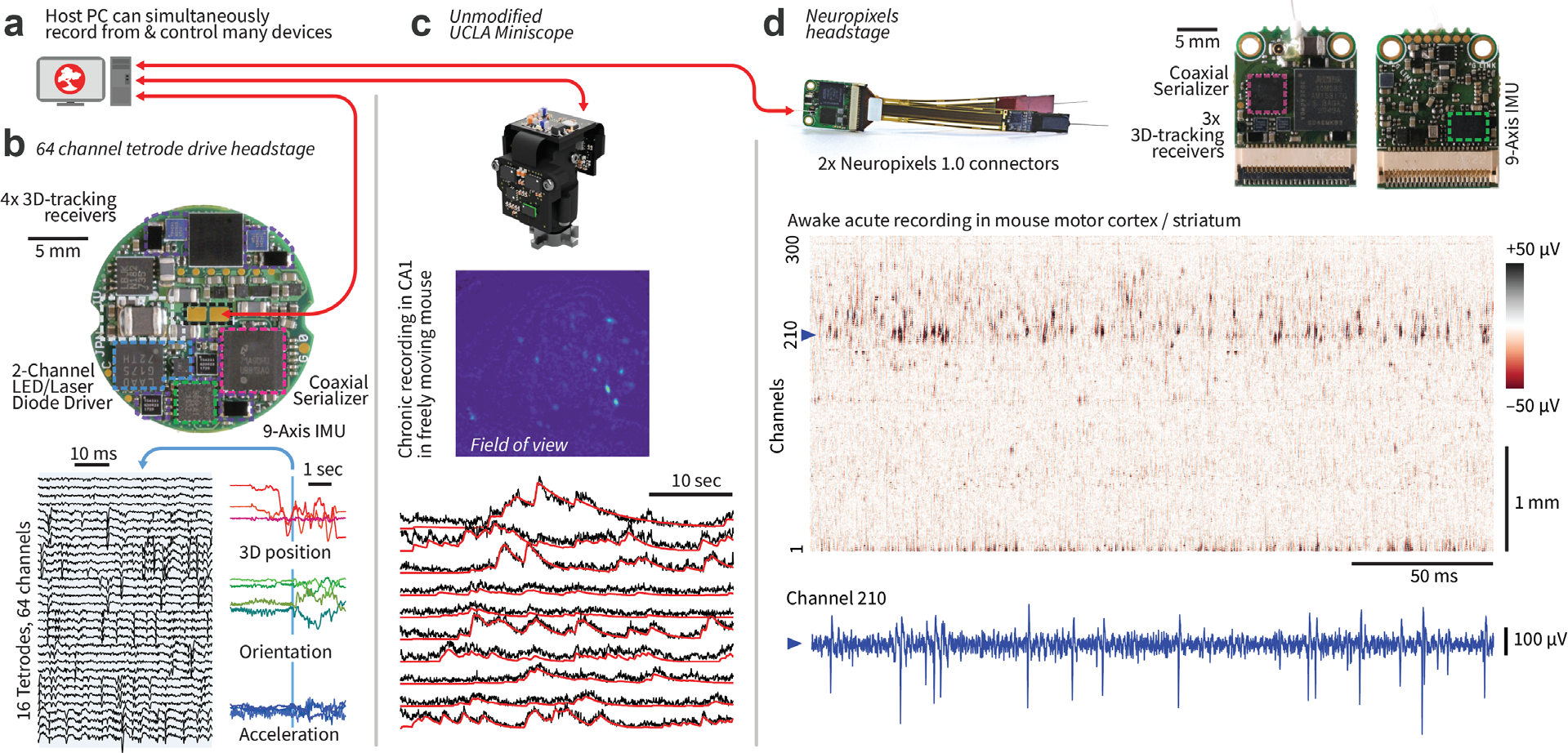
ONIX is compatible with existing and future recording technologies. **a**, ONIX, together with Bonsai, can simultaneously record from and synchronize multiple data sources, such as tetrode headstages, Neuropixels headstages, and/or UCLA Miniscopes. **b**, Top: a 64 channel extracellular headstage, as used in Figs.1-3, with 3D-tracking, electrical stimulator, dual-channel LED driver, and IMU (bottom side; not shown). Bottom: example neural recording and corresponding 3D pose traces collected from the headstage. **c**, ONIX is compatible with existing UCLA Miniscopes (versions 3 and 4)^45,55^. Middle: Maximum projection after background removal of an example recording in mouse CA1. Bottom: Background-corrected fluorescence traces (black) and CNMF output (via Minian^57^, red) of 10 example neurons. **d**, An ONIX headstage for use with 2 Neuropixels probes, and IMU to enable torque-free commutator use for long-term freely behaving recordings. A voltage heat map shows a segment from a head-fixed recording. A voltage time-series from the channel indicated by the dotted line is shown in blue.

ONIX is capable of sub-millisecond closed-loop neural stimulation on a standard (non real-time) operating system (Fig. 1e, Suppl. Figs.4 & 6). This level of latency is otherwise only achievable on specialized operating systems^52^ or hardware^53^. Further, its hardware-agnostic, open-source C API allows scientists to more easily develop, share, and replicate algorithms for such experiments in commonly used programming languages such as C++, C#, Bonsai^54^, Python, etc., without being tied to specific hardware.

To demonstrate ONIX’s ability to perform rich, uninterrupted studies of freely moving mice, we performed ∼8-hour recordings while mice explored a 3D arena. The arena was a 1.5 × 1.5 × 0.5 m arena constructed from hexagonal blocks of styrofoam, cut to different heights, giving mice the opportunity to run, climb, and jump (Figs. 2,3, Supplementary video 1). We exposed naïve animals implanted with micro-drives^44^ to this unfamiliar environment without behavioral shaping or human intervention. We compared the mouse behavior achievable with the ONIX system to a typical, modern acquisition system (Fig. 2c-e), by attaching a standard tether (Intan “SPI Cable”, 1.8 mm diameter), counterweighted with an elastic band to eliminate tether weight, alongside the micro-tether. This allowed use of the ONIX headstage for position measurements and commutation/untwisting, effectively adding torque-free commutation to the SPI tether, while imposing the mechanical effect of the weight of a traditional tether on the mouse. The tethers were switched every 2 hours over an 8-hour recording session (Fig. 2e). Except for tether exchanges, no experimenter was present. Even with our zero-torque commutation, the additional forces imposed by the standard tether (head-torque >0.4 mNm, measured in separate experiment, Fig. 1d) had a substantial deleterious effect on exploratory behavior and freedom of head movement (Fig. 2c-e, quantified via entropy of the spatial occupancy and head position distributions, median entropy 4.21 vs. 0.287 bits for spatial occupancy, and 0.75 vs. 0.50 bits for heading, P<0.0001, Wilcoxon rank sum test, see methods). When only the micro-coax tether was used (head-torque ≈ 0.1 mNm), the animal resumed free exploration of the arena (Fig. 2e). Some digital tethers, for example the twisted pairs used with Neuropixels^43^, would fall in between the standard SPI cables used here and the ONIX micro-coax in terms of weight and flexibility, but they would lack zero-torque commutation, impacting behavior and limiting recording duration.

To compare the behavior achieved with ONIX to an implant-free condition, we used 5 synchronized cameras for marker-less 3D tracking^31^ of non-implanted mice (Fig. 2e, left). The degree of arena exploration and the head orientation distributions of ONIX vs. non-implanted mice were statistically indistinguishable with 4 hours of data per condition yielding largely overlapping confidence intervals (one mouse per experiment, quantified via entropy of the occupancy distributions, 95% confidence intervals for spatial occupancy: [0.208,0.369] non-implanted vs. [0.224,0.383] ONIX, and confidence intervals for heading: [0.479, 0.536] non-implanted vs. [0.474, 0.533] ONIX, see methods). The median and maximum running speed of the implanted mice was reduced by a factor of ∼2 compared to animals with no implant. However, the ONIX micro-coax provided a ∼12x increase in median running speed and a ∼2x maximum speed compared to the standard tether (Fig. 2d).

To demonstrate the utility of long recordings without behavioral disruption, we conducted a 7.3-hour recording with a tetrode drive implant^44^ in retrospenial cortex in the 3D arena (Fig. 3). Mice spontaneously jumped to heights of >10cm (Fig. 3a,b), allowing us to observe neural activity during jumps (Fig. 3f). This behavior was absent in mice with the heavier tether.

Finally, we demonstrate the ONI standard’s flexibility by using the same ONIX system to acquire from and control two additional widely-used, third-party devices: UCLA Miniscopes^45,55^ (Fig. 4c, Suppl. Fig. 5) and Neuropixels probes^43^ (Fig. 4d). These recordings can also be performed simultaneously if needed. By using the Bonsai software^54^ for data acquisition, we also demonstrate integration of synchronized multi-camera tracking (Fig. 2, Suppl. Fig. 5). Bonsai enables the integration of real-time processing tools such as animal tracking via real-time DLC^56^ or SLEAP^24^, enabling experiments that react to animal behavior with high precision.

For developers, the ONI hardware standard and API streamlines the development of new probe and sensor technologies into headstages that have immediate integration with existing technologies, which lowers the barrier for individual labs to create custom instruments designed for specific experiments. Similarly, ONI simplifies the development of completely new data acquisition systems by providing a scalable, easy-to-use interface for communication between software and FPGA firmware, and ensures interoperability between these systems.

In sum, our system provides a probe-agnostic open-source interface for use in neuroscience. It allows, for the first time, long and high-bandwidth recordings in mice and similarly sized animals during naturalistic behaviors comparable to those of non-implanted animals. This ability will accelerate progress in many areas of research, such as motor learning^58^, sensory processing during natural behaviors^8^, social behaviors^59^, play^9^, or on cognitive aspects of spatial behaviors^60,3,61,62^ by allowing unimpaired motor behavior, reducing animal fatigue over time, and enabling navigation in large environments.

## Code and design file availability

The full ONI specification is available at https://github.com/open-ephys/ONI. Design documents for the described ONIX hardware are available as follows -- Host interface: https://github.com/open-ephys/onix-fmc-host, Breakout board: https://github.com/open-ephys/onix-breakout, 64-Channel Intan headstage: https://github.com/open-ephys/onix-headstage-64, and Neuropixels headstage: https://github.com/open-ephys/onix-headstage-neuropix1. Software along with extensive hardware and API documentation is available at https://open-ephys.github.io/onix-docs. Firmware can be made available upon request as configurable IP blocks. Unless specified in the respective repositories, all material is distributed under the creative commons CC BY-NC-SA 4.0 license and is therefore free to adapt and to share with appropriate attribution and under the same license for non-commercial purposes.

## Acknowledgements

We thank the Open Ephys Production Site team for beta-testing, Emily J. Dennis, Antonin Blot for additional system testing, Joaquim Alves da Silva and Hanne Stensola for testing of the 3-D tracking system, Andrew Bahle for help with testing the Neuropixels headstage. Albert Lee and Emily J. Dennis for comments on the manuscript. Funding Sources - Jonathan P. Newman: National Institute of General Medical Sciences T32GM007753 (E.H.S.T), the Center for Brains, Minds and Machines (CBMM) at MIT, funded by NSF STC award CCF-1231216, and NIH 1R44NS127725-01 to Open Ephys Inc. Jie Zhang: NIH 1R21EY028381. Takato Honda: Picower Fellowship, JPB Foundation and Picower Institute at MIT, Osamu Hayaishi Memorial Scholarship for Study Abroad, Uehara Memorial Foundation Overseas Fellowship, and Japan Society for the Promotion of Science (JSPS) Overseas Fellowship. Marie-Sophie H. van der Goes: Mathworks Graduate Fellowship. Mark T Harnett: NIH R01NS106031 and R21NS103098, was a Klingenstein-Simons Fellow, a Vallee Foundation Scholar, and a McKnight Scholar. Matthew A. Wilson: National Science Foundation STC award CCF-1231216, and NIH TR01-GM10498. Jakob Voigts: NIH 1K99NS118112-01 and Simons Center for the Social Brain at MIT postdoctoral fellowship.

## Contributions

ONI concept: JPN; ONI spec and API: ACL, JPN, GL; Firmware prototype: JZ, JPN; Firmware implementation: JPN, JZ, ACL; Headstage hardware development: JPN, JZ, ACL; Headstage beta-testing: MSHvdG, NJM, TH; Commutator: JV, JPN; Mouse electrophysiology: JV, NJM; Neuropixels experiments: JHS, AL; Behavior comparison experiments: JV, NJM, JPN; Data analysis: JPN, JV, JS; Miniscope recordings: TH, JZ; Documentation: JPN, ACL, AHL; Beta-testing coordination: AHL; Manuscript: JV, JPN, MTH, MAW, with input from all authors.

