## Supplementary Figures and Methods for "A unified open-source platform for multimodal neural recording and perturbation during naturalistic behavior"

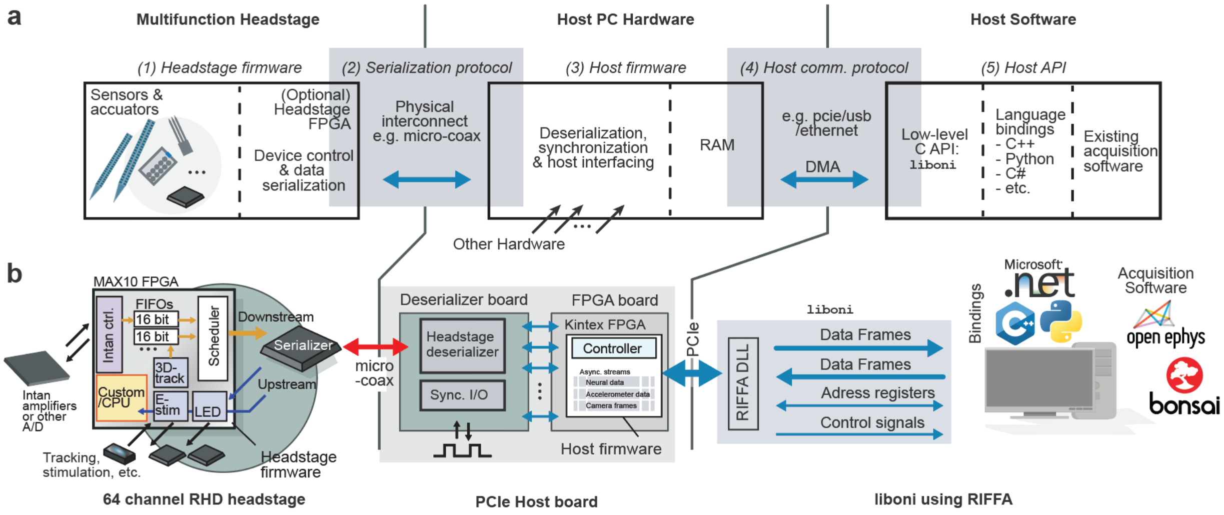


**Supplementary Figure 1: ONIX system block diagram**. **a,** The ONIX system architecture consists of 5 building blocks: (1) headstage firmware, (2) data serialization protocol, (3) host firmware, (4) host communication protocol, and (5) application programming interface (API). The firmware modules and host API (open black boxes) are both specified and implemented within this project. These are reusable across host hardware and physical interfaces (e.g. USB, ethernet, PCIe, etc). The serialization and host communication protocols (grey boxes) are generically specified by ONI. This project focuses on a micro-coaxial serializer to implement this spec. Other communication options would require custom, ONI-conforming firmware to be written. **b,** Each element of the architecture in panel (**a**) applied to the 64-channel Intan headstage used in Figs 1-3. The serialization protocol and host communication protocol are implemented using a micro-coaxial serialization link and PCIe, respectively.

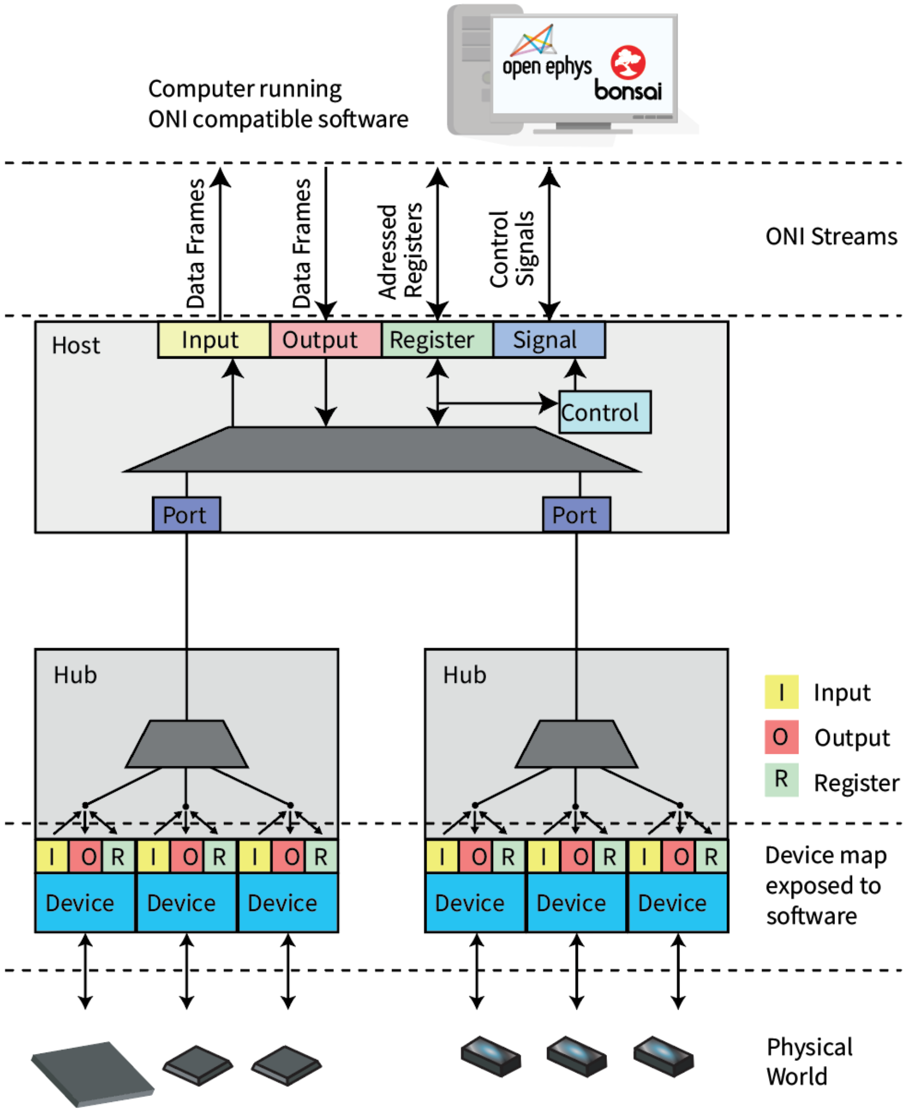


**Supplementary Figure 2**: ONI **communication block diagram.** Hubs (e.g. headstages, miniscopes, etc.) aggregate data and control sensors and actuators that interact with the world. Each sensor is provided with a separate, low-latency FIFO and control block. Configuration commands are demultiplexed from the host communication link to control each device on the headstage. Sensor data is multiplexed into a shared high-bandwidth link by a multiplexer on each hub. Low-bandwidth bidirectional configuration data is also transmitted over this link. In the hardware described in this paper, this link is implemented over a micro-coaxial serialization link, but could also be implemented using other wired or wireless links. Sensor data is demultiplexed by the host serial interface to create a local representation of the hub. This allows transparent control of each hub from the host computer, and low-level communication with the physical headstage via the serialized link is hidden by the firmware. Sensor and configuration data are streamed to and from the host PC using a high-speed bus, such as PCIe or USB3.0. The same API is used regardless of the physical communication interface. The appropriate drivers and translation layers are dynamically loaded for each physical interface that is used.

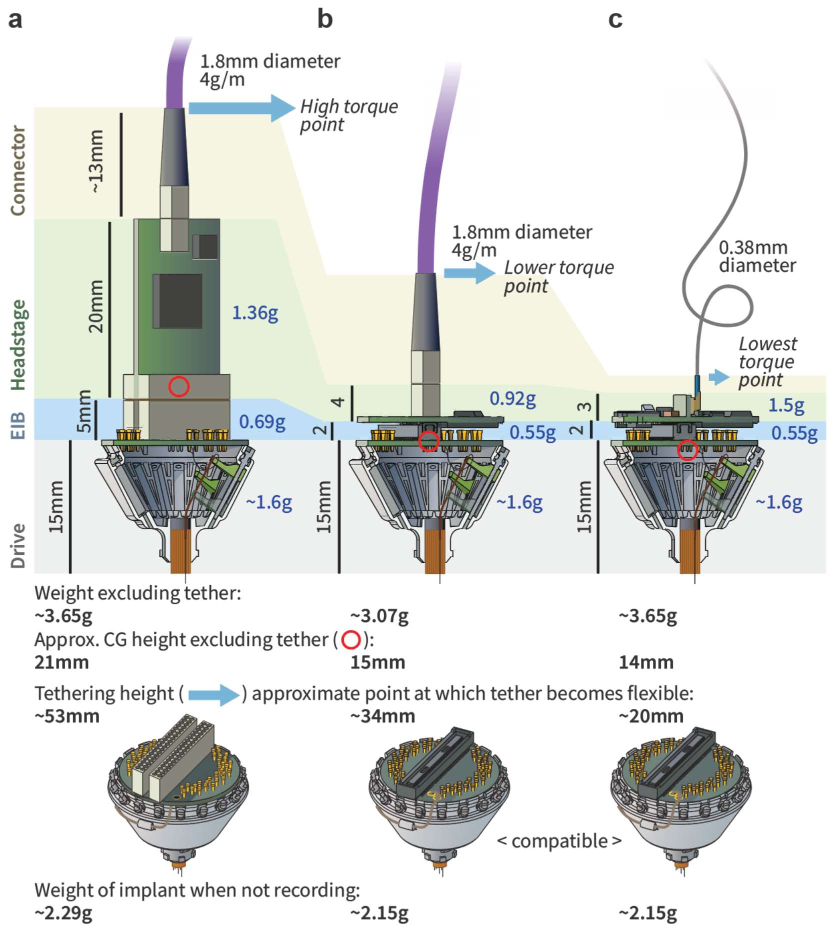


**Supplementary Figure 3. Relative weights and torque arms of comparable headstages.** Comparisons are between headstages on the same low-profile 16 tetrode drive implant[44]. **a,** Current Intan-based headstage or comparable system. Omnetics connectors, vertical headstage layout, and tether all contribute to height and long moment arm, so that sideways forces on the tether exert strong rotational forces on the animal's head. **b,** Best-possible low-profile headstage with current digital tethers, assuming a single, flat PCB design equivalent to the ONIX headstages described here. The digital tether still leads to a significant moment arm. **c,** In addition to a low-profile headstage, the ONIX micro-coax further lowers the torque arm, and also practically eliminates torque loads by virtue of an extremely light and flexible tether.

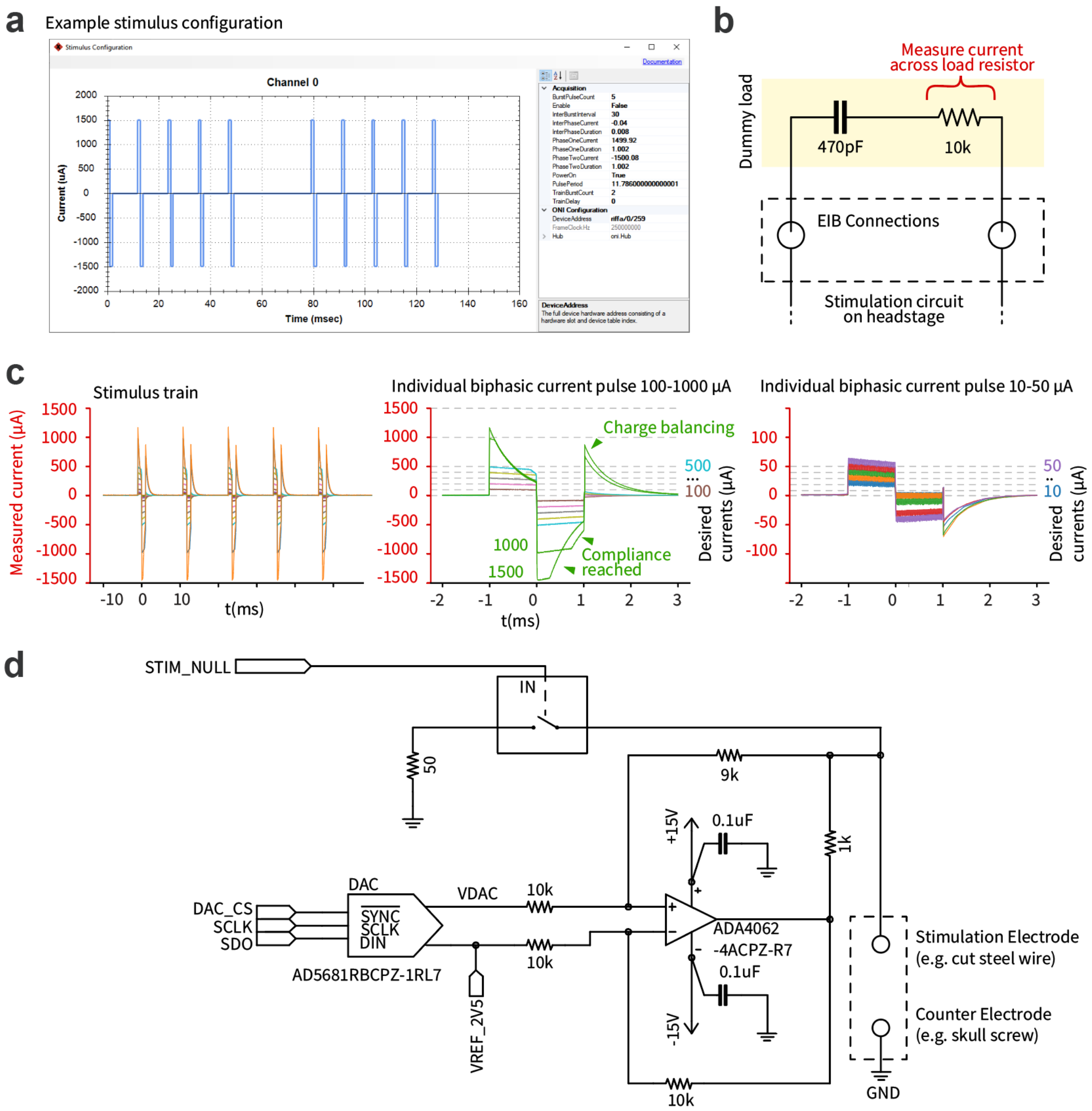


**Supplementary Figure 4.** Integrated electrical microstimulation circuit on ONIX's 64-channel Intan headstage. **a,** example pulse train configuration via graphical user interface in Bonsai. **b,** Setup for characterizing the circuit: The stimulator was used to drive biphasic current pulses into a model load (47 nF capacitor in series with 10kOhm resistor to ground), representing a high impedance stimulation scenario. The voltage across the 10 kOhm resistor was measured with an oscilloscope to monitor the delivered current. Pulse trains consisted of 1 ms per phase, biphasic, positive first pulses with inter-pulse interval of 10 msec (83 Hz pulse to pulse period). **c,** Measurement results: the amplitude of both phases was varied from ±10 uA to ±1.5mA. For pulse-based stimulation, each phase must satisfy the following inequality to be achievable with the system's maximum voltage: $|I_{stim} * (R_{electrode} + t_{phase}/C_{electrode})| <= 15V$. In our test setup, for ±500 uA, this results in ~15.6V per phase which is why voltage compliance was reached for this target current. If the compliance voltage is reached, or pulses are delivered with asymmetric positive and negative charges, a charge imbalance will occur, resulting in a residual electrode voltage following the conclusion of the stimulus. The large artifacts following such failed stimuli result from the onboard charge balancing circuit removing this voltage after each stimulation pulse concludes. The mild slopes on the successfully delivered pulses are due to the finite voltage slew rate of the stimulation circuit. The stimulus current is recorded though an auxiliary channel on the Intan chip for verification of stimulus current waveforms. **d,** Simplified electrical stimulation circuit. A DAC controls an improved Howland current pump. Electrodes are discharged in between pulses to reduce artifacts and impose charge balancing. Control signals provided by the headstage FPGA are shown to the left. Elements are referred to in the text using their labels.

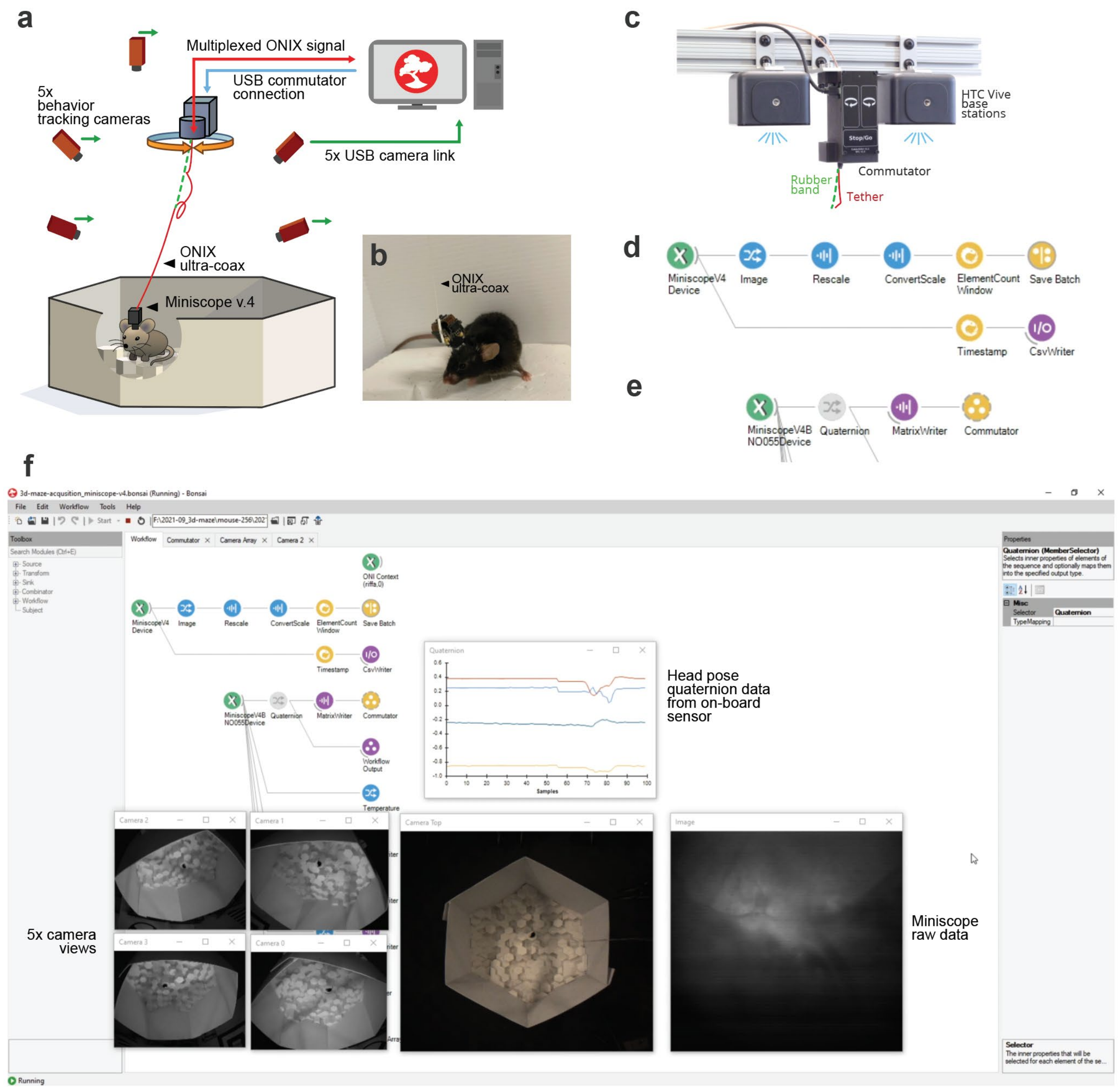


**Supplementary Figure 5. Example setup and Bonsai workflow for simultaneous neural recording and behavior tracking. a,** A mouse with a Miniscope implant is free to explore the same arena described in Figs.2&3, and is being tracked by 4 side-mounted, and one overhead camera. The entire experiment is performed using the Bonsai[54] software. In addition to the headstage connection (red), Bonsai controls the motorized commutator (blue) via a serial-over-USB link, as well as the 5 cameras (green) using the camera vendor API over USB. **b,** Picture of a mouse with a Miniscope v.4 implant[45], controlled by Bonsai via the ONIX system. **c,** Close-up of the motorized commutator and 3D-tracking base stations. **d,** Excerpt of the Bonsai workflow used to record the Miniscope image stream. Data arrives seamlessly via the 'MiniscopeV4' node (green) in Bonsai, which communicates with a standard Miniscope through the ONIX micro-coax, is then rescaled using common Bonsai image processing nodes (blue) and saved to disk (yellow). A separate data path assigns a time stamp to each Miniscope frame, allowing data to be synchronized with behavior cameras or other data that were not acquired synchronously. **e,** Part of the Bonsai workflow used to drive the torque-free commutator (Fig.1b). The head orientation quaternion output of the 'MiniscopeV4 BNO55' node (green) is one of the possible outputs of the BNO55 chip on the headstage (the same chip is shared across the Miniscope, and the 64 channel Intan, as well as Neuropixel headstages described here), in addition to other outputs such as acceleration, temperature, and gravity vector (not shown here). The quaternion is the saved to disk as raw data (purple), to be used with an associated timestamp (not shown here, same procedure as for the Miniscope timestamps in panel d), and sent to the 'Commutator' node (yellow), which drives the commutator to follow the animal's rotation and remove any twisting of the tether. **f,** Example screenshot from Bonsai showing simultaneous data acquisition from all sources. Example workflows are available on the ONIX GitHub repository (https://open-ephys.github.io/onix-docs/Software%20Guide/Bonsai.ONIX/Bonsai%20Examples/index.html).

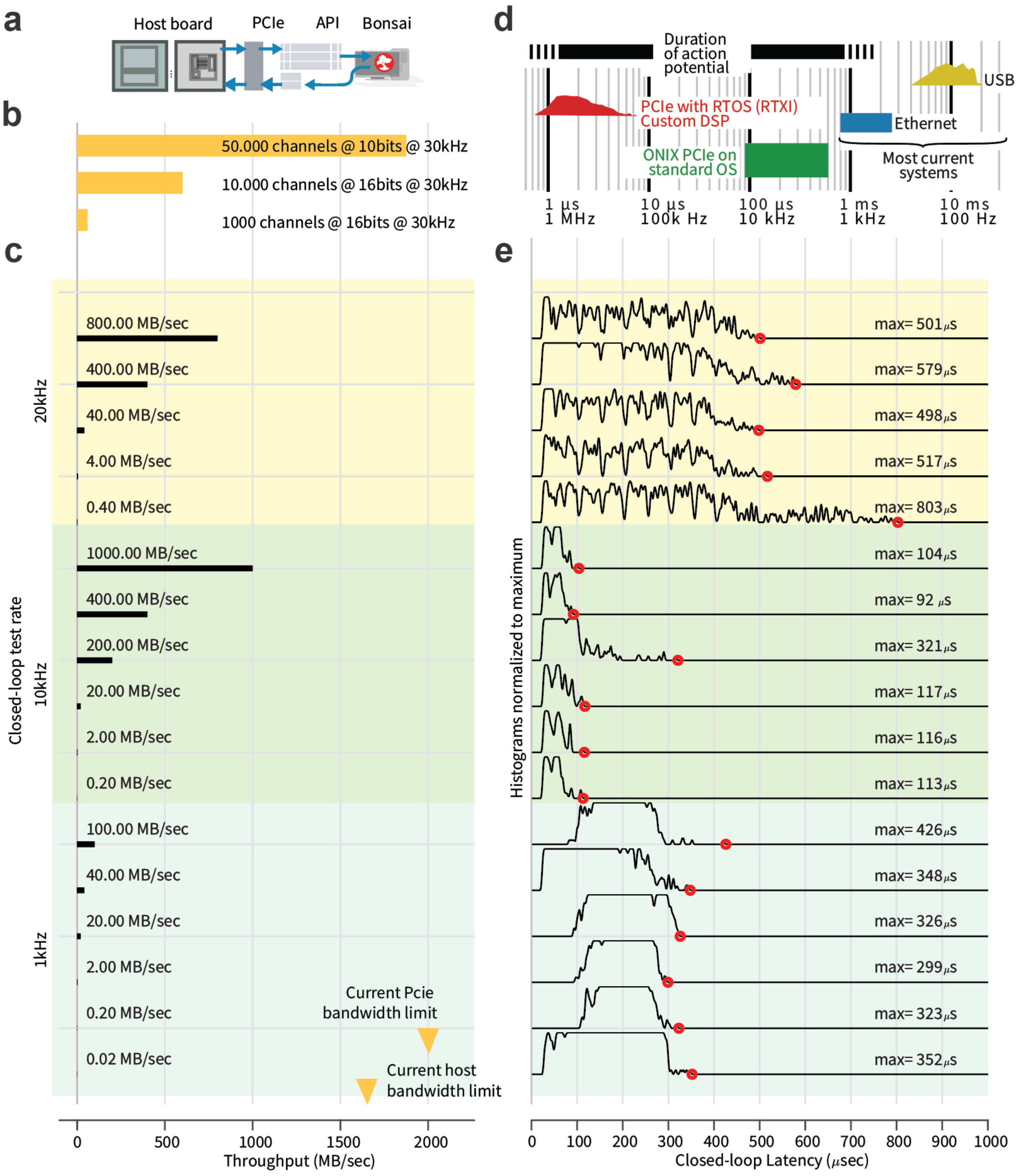


**Supplementary Figure 6. Host interface latency.** Latencies were measured for a closed-loop round-trip from the PCIe host to C software and back. **b,** Example theoretical data rate requirements for some high channel count recordings at 16 or 10 bit sample depth. Actual data rate requirements may differ depending on communication overhead. Future work could reduce the bandwidth requirements by compressing data on the headstage[63,64]. **c,** Different frequencies at which the closed-loop tests were run, each with increasing buffer sizes and resulting theoretical bandwidth. Note that the actual bandwidths can be limited by the headstage interconnect. **d,** Overview of latency ranges achievable with other technologies. **e,** Distribution of measured round-trip latency for all settings on a Windows 10 desktop computer with an Intel i5 CPU. The lowest latencies were measured with a 10 kHz rate. Results will differ on different hardware/OS, and depend on system load.

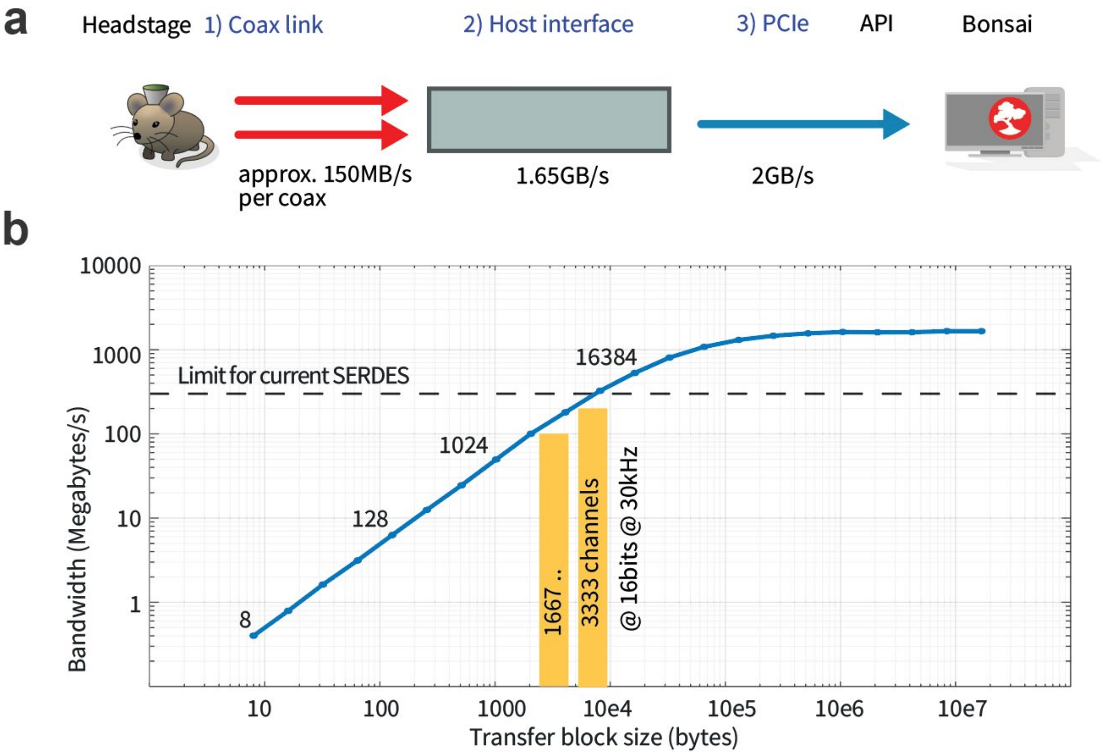


**Supplementary Figure 7. Real-world data rates of the ONIX system. a,** Overview of the system bandwidths. Bandwidth limits apply to different parts of the system: 1) The current serial over coax (Serializer-deserializer, SERDES) chip used on the headstages described here (Texas Instruments FPD-Link III for r 1-MP/60-fps), using a 100MHz clock rate on a 12-bit interface, results in 1200Mbps (150MBps) bandwidth. The currently highest used bandwidth is ~48MBps on the Neuropixels headstage. Future implementation with other SERDES chips can improve this bandwidth, with the current headstage deserializers imposing a limit of ~300MBps. 2) The host interface currently runs an internal 64bit bus at 250MHz and is capable of 1.65GB/s. 3) Finally, the 4-lane PCIe interface used by ONIX is theoretically capable of ~2GB/s, of which we can currently use 1.65GB/s due to protocol overhead. It would be trivial to expand to higher lane counts in the future. **b,** To measure real-world throughput, the ONIX host has a load testing FPGA-core with a configurable packet size and rate, which result in a given bandwidth. For a set of given transfer block sizes (here, powers of 2 starting at 64 bytes were used), we gradually increased the load bandwidth until the output FIFO, which is implemented on external RAM, started to fill, indicating that the transfer speed couldn't cope with the data production. The resulting curve therefore corresponds to empirically achievable data rates. Measured bandwidth from data generated in a test device as a function of transfer block size. Yellow: Approximate data rate requirement for example numbers of electrophysiology channels, sampled at 30KHz and 16 bit with no optimization.

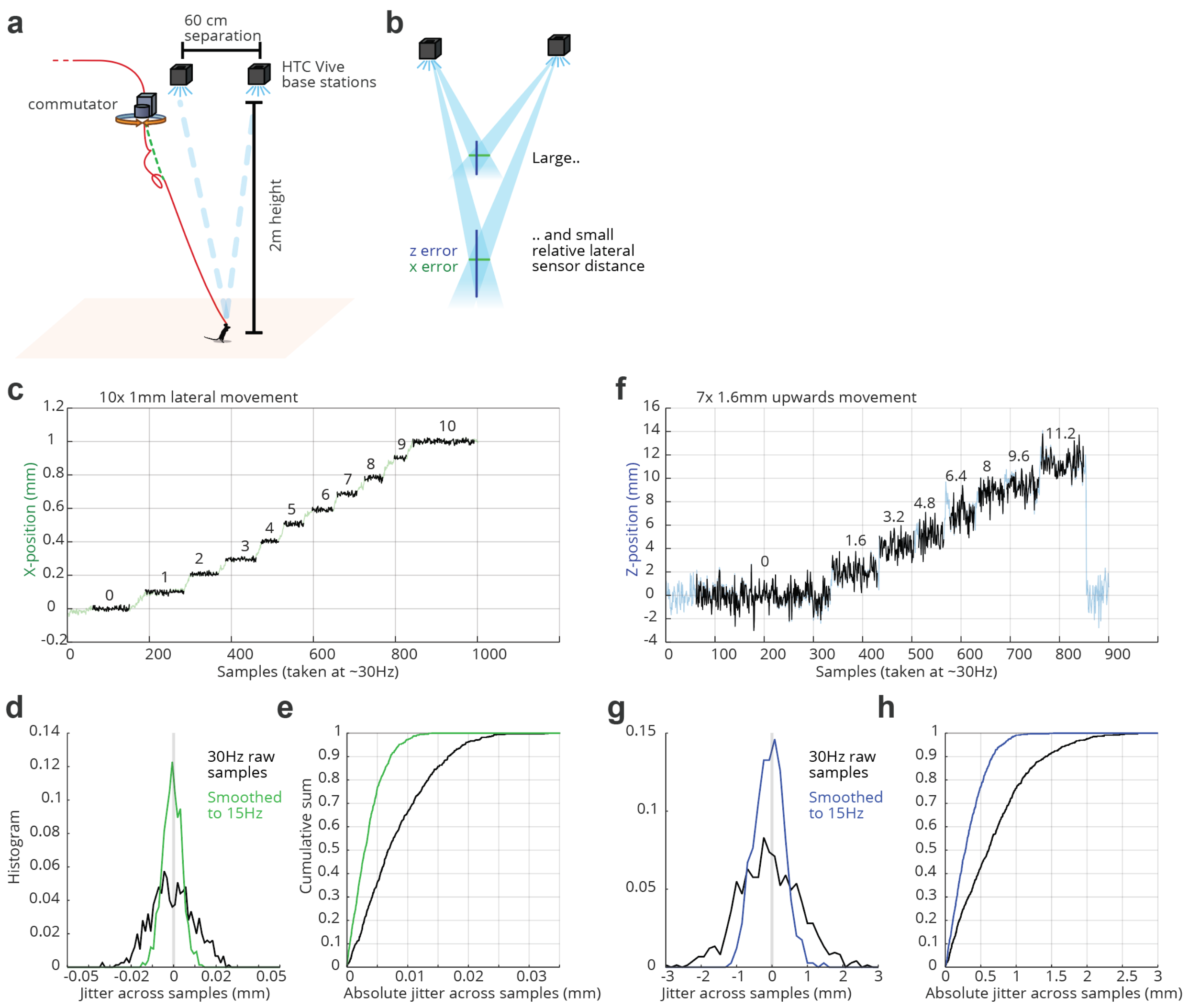


**Supplementary Figure 8**. **Precision of the 3D-tracking. a,** Experimental setup: A ONIX headstage was moved by known amounts at a 2-meter distance from the base stations. The base station pair was separated by 60 cm. **b,** Coordinates are derived from angular tracking data relative to the known base station positions. In our case (60 cm separation at a 2 m distance), this results in worse resolution in the height/Z direction than in the lateral/XY direction. **c,** Raw position measurements for known step displacements. **d,** Measurement jitter relative to known positions. In addition to the raw sample data, smoothed data at 15 Hz are plotted. **e,** Resulting cumulative distribution of absolute jitter. **f-g**, same as c-e but for height/Z measurements, this setup with large distance from the base stations and small distance between the base stations demonstrates degraded Z-resolution. Higher Z resolution can be achieved at the same distances by increasing the spacing between the base stations above 60 cm.

### Supplementary methods:

The following methods section provides a high-level overview of the methods, experiments, and protocols described in this manuscript. For in-depth and up-to-date documentation of the hardware, software, and detailed user guides, consult the documentation at the appropriate repository on https://github.com/open-ephys, and the associated higher-level documentation at https://open-ephys.github.io/onix-docs.

#### Design Goals and High-Level Architecture

Neuroscience data acquisition systems often use proprietary interfaces and protocols for device communication in the name of performance and commercial advantage[65]. Some side effects of this situation are device lock-in and difficulty of device interoperability. On the other hand, while open-source hardware, including our own designs, provides access to communication protocols and hardware, these protocols and physical interfaces generally underperform commercial options and tend be highly non-standard and brittle, making them difficult to extend and maintain.

Table 1 summarizes physical interfaces, firmware, communication protocols, and host APIs used by several open source hardware projects in widespread use for systems neuroscience research. Each of these projects employs a different set of interconnects, device drivers, firmware, and APIs. Their designs are tailored for the features of a particular sensor and are difficult to extend to general purpose data acquisition. Further, each of these solutions requires linear scaling of headstage tether conductors with the number of sensors at the headstage, and precludes the combination of high-throughput recordings with naturalistic behavior.

| | **Intan / 'Classic' Open Ephys** | **Neuropixels (Ver. 1.0)** | **UCLA Miniscope** | **ONIX** |
|---|---|---|---|---|
| **Headstage** | | | | |
| Connection to host | SPI via LVDS twisted pairs | Coax or twisted pair | Coax or twisted pair | Coax |
| Onboard processing | None | None | None | FPGA |
| Connector scaling | 8 additional conductors per chip | Limited to one probe | Limited to single camera sensor | Limited by Bandwidth (1GPps equiv.) |
| Arbitrary chip support | No | No | No | Yes |
| **Host Interface** | | | | |
| Connection to PC | USB | PXI, PCIe | USB | PCIe |
| Communication protocol | Opal-Kelly Front-Panel + Drivers | Ad-hoc serialization | USB-webcam | Probe-agnostic serialization |
| Programming interface | Rhythm API (Intan chips only ) | NeuroPixels-specific API | Cypress SDK for webcams | Probe-agnostic ONI API |
| Firmware | Rhythm Verilog (Intan chips only) | Neuropixels-specific | Cypress Firmware | ONI IP block + arbitrary probe-specific firmware |
| **Software Support** | Open Ephys GUI, Bonsai, Intan RHX | Open Ephys GUI, SpikeGLX | Miniscope GUI, Bonsai | Open Ephys GUI, Bonsai, ONIX console app |
| **Closed Loop Latency** | 10s of milliseconds | ~3 milliseconds | 10s of milliseconds | <100 microseconds |
| **Compatibility Limitation** | Only supports Intan Chips | Only supports Neuropixels | Only supports camera sensors | Compatible with Intan chips, Camera sensors, Neuropixels, and any other sensors |

**Table 1:** Summary of communication data-acquisition architectures, host interfaces, software-support, closed-loop performance, and notable limitations for three widely used open source hardware projects for in-vivo electrophysiology.

To address these limitations, we have taken a top-down approach to design communication protocols, firmware, and host API to support acquisition from any mixture of devices (Fig. 1a, Fig4). Specifically, our designs meet the following requirements:

1. Heterogeneous device acquisition and control
   a. Well-defined communication protocols that support bidirectional communication with any mixture of sensors or actuators
   b. Generic, low-level API that takes inspiration from existing, widely-used libraries and is focused on the creation of high-level language bindings.
   c. Bus-agnostic: firmware, protocols, and API should not rely on a particular physical layer between headstage and host computer (e.g. wireless or tethered) or host interface and CPU (e.g. USB, Ethernet, or PCIe).
2. Closed-loop performance
   a. Sub-millisecond headstage to host-PC round trip time
   b. Ability to drive neural stimulation devices directly from headstage
   c. First-class bidirectional communication with host computer
3. Practical in-vivo use
   a. On-headstage processing
   b. Generic data serialization protocol and firmware can be used with single-wire tether or wireless transmission
   c. Physical headstage connectorization bounded to a single coaxial connection regardless of the number of devices on heastage
   d. Bandwidth for up to 1000s of recording channels sampled at 30 kHz
4. Use standard parts and interfaces
   a. Make use of standard parts created for large markets (mobile computing, automotive, etc)
   b. Use standard and widely used connectorization, and avoid specialty hardware wherever possible
   c. Use standard UNIX programming practices with minimal dependencies
   d. Cross platform
   e. Minimize cost

The resulting system consists of 5 major components (Suppl. Fig. 1). Reusable headstage firmware modules (Suppl. Fig. 1 (1)) allow headstages to interface with an arbitrary arrangement of sensors and actuators. Device data streams are combined and packaged at the headstage using generic and extensible firmware modules and a well-defined serialization protocol (Suppl. Fig. 1 (2)) over a micro-coaxial cable (serialization protocol is interconnect-agnostic and could use a wireless link). Data are deserialized by the firmware running on a FPGA interface board inside the host computer (Suppl. Fig. 1 (3)). A device driver works with firmware running on the deserialization hardware to pass data to user-space memory using a high-speed communication protocol developed to support communication with arbitrary remote and/or local data sources and sinks (Suppl. Fig. 1 (4)). Any peripheral bus capable of realizing this protocol is supported, but we focus on use of the PCIe bus due to its high performance. Finally, a low-level C API (Suppl. Fig. 1 (5)) provides software access to resultant data and control streams. This API's simplicity makes it cross-platform and amenable to the creation of language bindings for easy integration with existing acquisition software. This architecture greatly decreases and stabilizes closed-loop response latencies compared to existing open-source systems and permits scalable combination of miniature head-borne sensors and actuators to simplify complex closed-loop experimentation.

#### Open Neuro Interface (ONI): A standard for head-borne instrumentation

This work was built on the ONI specification, a set of general-purpose communication protocols, device driver specifications, and programming interfaces to support arbitrary mixtures of hardware. The ONIX system outlined here is one implementation of this specification.

Open Neuro Interface (ONI) is an open standard describing a high-speed interface between a computer and a collection of devices, which can be of different nature. Its goal is to provide a single, unified protocol to communicate with the variety of instruments widely used in neuroscience such as electrophysiology acquisition devices, tracking systems, cameras or stimulators. It defines both a logical structure for the different hardware elements as well as streaming and configuration protocols, along with API specifications. It, however, does not define a specific hardware transport layer, leaving that open to different implementations, but only how the data should be organized. The specification itself can be found at https://github.com/open-ephys/ONI and provides a comprehensive documentation. Suppl. Fig. 2 provides a simplified overview of the structure of the standard.

#### Acquisition performance

Performance of any acquisition system is measured in bandwidth, indicating the amount of data it can acquire per second, and latency, which measures the time between an actual event and the ability to act on it. The maximum bandwidth allowed by the x4 PCIe Gen2 interface is 2GB/s. However, the current ONIX system operates at a maximum theoretical bandwidth of 1.6GB/s total. Data transfer is not realized continuously, but in blocks, with each individual transfer operation featuring a small overhead. As such, bigger block size increases actual bandwidth by reducing overheads, at the cost of increased data latency. Suppl. Fig. 6 shows measured bandwidth using a load-testing device in the host firmware.

While latency, measured in time, is related to block size, its actual impact is dependent on the data origin. This effect is more detrimental for devices generating small loads at high frequencies, which are rarely the case in neuroscience, as opposed to big sample sizes at the KHz range. For example, a 8192-byte block size would introduce a latency of 1024 samples on a simple, 8-byte timestamping device. However, for the case of 4 neuropixel probes, with a sample size of 480bytes each, a 8192-byte block transfer would imply a latency of less than 5 samples. Although total latency is dependent on block size, there is a fixed, minimum latency associated to the processes of transfer initiation. For the ONIX system, transmission latency was measured with the smallest block size and a simple C program responding to a digital event. Under these conditions, 150µs of maximum transmission latency were measured. Figure 1e shows closed-loop latencies measured for a round-trip from electrophysiology data to optical or electrical stimulation on the headstage with a 64 channel intan-headstage.

#### Headstages

Using the architecture detailed above, a headstage for extracellular electrophysiology, using amplifier / digitizer chips by Intan tech (https://intantech.com/) was designed for use in rodents, and one headstage for use with two neuropixels[43,66] probes. Images of these headstages and listings of major features are provided in Figure 4. In the following text we provide detailed descriptions of each of these features.

##### Size and Weight

The Intan headstage is designed to be mounted flat on top of the electrode interface board (EIB) of an tetrode microdrive (see our companion paper for drive designs[44]). This parallel, rather than the traditional orthogonal mounting scheme was chosen to reduce torque on the animals' head during freely moving behavior (Suppl. Fig.3). Headstage-64 is 19-mm in diameter. When PCB is completely populated, it weighs ~0.95 grams. The neuropixels headstage was designed as a traditional dual-probe layout.

##### Headstage IO, Serialization, and Physical Interface

A key feature of the firmware provided with the project is the ability to serialize an arbitrary set of asynchronous data sources into a single data stream. This stream can then be transmitted back to the host using any transceiver with appropriate bandwidth. Our choice of transceiver between our headstage and the host PC was inspired by the UCLA miniscope, which uses a Texas Instruments DS90UB913A/14A serializer/deserializer (SERDES) pair to combine power, high-bandwidth data transmission, and low-bandwidth headstage control into a single microcoaxial cable. Although our headstages focus on the use of this transceiver, it is important to note that the modular design of our firmware allows any appropriate link to be used. For instance, wireless communication, e.g. via wifi, or Bluetooth, could also be used to enable real-time data streaming.

In the tethered configuration, the coaxial cable is the only external connection to the headstage. Power (DC), a control "back-channel" (70 MHz), and a data "forward-channel" (700 MHz) occupy different portions of the RF spectrum and therefore can be resolved as distinct signal streams. Power is DC-coupled out of coaxial interface via an inductive path while a capacitive path from the serializer is used to AC couple in the control and data signals. The reverse occurs at the deserializer. Although this chip is intended for use with camera sensors, it can be repurposed to transmit arbitrary data using the headstage and host FPGAs to provide a camera-like digital interface for arbitrary data. We do this by repurposing the SERDES data and control interface using an FPGA to spoof a camera and camera decoder at serialization and deserialization ends, respectively.

##### Headstage FPGA

The current 64 channel and neuropixels headstages use the Intel MAX10 FPGAs for peripheral device control (e.g. stimulation timing), sensor sample collection, data packaging and buffering, and serializer interfacing on the headstage. This FPGA was chosen due to its small size, integrated flash storage, PLLs, and 32-bit soft processor. The 64-channel headstage uses an 81-pin wafer level chip scale package (4x4 mm footprint, Fig. 1a, Fig. 4b).

#### Headstage Sensors and Actuators

##### Electrophysiology

The Intan headstage performs multichannel electrophysiology using a BGA-packaged Intan RHD2064 64-channel bioamplifier/digitizer chip. The firmware and API permit read and write access to the RHD2064's 64 control registers from the host PC. The neuropixels headstage interfaces with the Neuropixels chip using a similar MAX10 FPGA to the 64-channel headstage. It shares other digital logic blocks for 3D pose tracking with the 64 channel Intan headstage but omits the optogenetic and electrical stimulation circuits.

##### Electrical Microstimulator

The Intan headstage features a single constant-current, bipoloar electrical stimulation circuit (Suppl. Fig.4). Connections to the stimulation circuit are routed through the high-density connectors on the bottom of the headstage to the EIB where static or movable stimulation electrodes can be attached using standard methods. The stimulator is an improved Howland current pump with a bipolar 15 V supply. Stimulation current is measured on headstage and routed to an auxiliary input of the Intan chip. Outside of stimulus pulses, the circuit provides charge balancing by shorting the stimulation electrode to ground. Component values have been chosen to optimize circuit stability over a wide range of electrode impedances. Operation remains stable for macroelectrodes (e.g. low-impedance cut stainless-steel wire and microelectrodes up to 1 MOhm at 1kHz). The circuits can produce up to 1.5 mA of bipolar current within the bounds of its ±15V compliance voltage range. Although this circuit consists of multiple components, the firmware and API provide an abstract control interface that allows high-level configuration and stimulus timing of the entire circuit. In this context, its operation is very similar to a Master8 or PulsePal[67].

##### LED/Laser-Diode Driver

The Intan headstage provides 2 high-current LED/laser diode drivers for optogenetic stimulation (On Semi CAT4016). The maximal current is set over a wide (~10 mA to 800 mA) range via an external digital potentiometer. The optical power can then be adjusted linearly and synchronously across all channels within this range over 8-levels per diode load. Like the electrical stimulator, this sub-circuit is controlled as a single device with the API using parameters similar to a Master8 or PulsePal[67].

##### 3D Tracking System

Both the Intan headstage and the neuropixels headstage provide a set of sensors for precise, room-scale 6 degree-of-freedom (DOF) head tracking. An integrated 9-axis inertial measurement unit (IMU) provides kHz-scale measurements of head-pose and angular acceleration. The 64-channel headstages uses a Bosch BNO55 which provides low-frequency compass data to determine absolute bearing. A select set of IMU control registers are exposed through the firmware and API for calibration and to adjust data output type. For instance, the BNO55 allows on-chip sensor fusion to directly report pose in some acquisition modes. This fusion mode can be used to drive the motorized commutator and keep the tether from twisting when the animal rotates.
In addition to the IMU, each headstage has multiple light-to-digital transceivers to capture laser sweeps from "lighthouse" tracking stations. Fusion of IMU data with laser sweep timing information can be used to robustly deduce millimeter level 3D position within a ~10 $m^3$ environment (see Suppl. Fig. 8). Each of these devices is treated as a separate data source by the API, and raw data is streamed to the host computer for each sensor for sensor fusion. In the future, it is conceivable that these operations could be moved to the headstage FPGAs embedded softcore processor and the entire tracking system treated as a single abstract device that sends pre-calculated 6 DOF pose information.

##### Host board

Following the ONI specification, the ONIX host board (Fig.1a) was designed to aggregate data from different hubs and devices and interface with the computer through a high-speed PCIe bus. The Numato Nereid board with a Kintex-7 FPGA was used as the base of the host system. A FMC daughter card was made containing all custom

electronics for interfacing with external devices. In the future, host boards with other interfaces such as USB can be developed that will function interchangeably, though might strike a different balance between ease-of-use, throughput, and closed-loop latency.
The main components of the FMC board are two ds90ub934 deserializers that communicate through a coaxial link with different hubs. External connection to the deserializers is made via MMCX connectors. These devices require two different power inputs require, 3.3V for I/O lines to the FPGA and 1.8V for the core. Those are provided through 3.3V pins in the FMC connector, originating in the Nereid board, and a DC-DC Buck converter to efficiently derive the 1.8V lines. Power for the external devices, transmitted through coaxial cables, is derived from a 12V source also provided through the FMC connector. Instead of being converted to a fixed voltage, a combination of configurable step-down converters and digitally controlled potentiometers was used to control the link voltage for each port. This can compensate for larger cables that cause higher voltage drop or adjust for different devices. The host board also contains interfaces for general purpose analog and digital signals. It features 12 analog lines that can be, through digitally controlled analog switches, independently routed to either a multichannel ADC or DAC. The board does not feature direct digital lines, however, but 5 high-speed differential pairs, 2 output and 3 input, which can be used to interface with a variety of digital systems.
For extra synchronization capabilities, the host board contains 2 buffered highspeed clock inputs and 1 clock output, accessible through coaxial MMCX connectors. It also features an internal connector with 4 differential pairs of configurable direction, designed to connect multiple boards to work in a synchronized manner.

### Support Hardware

**Active motorized cable commutator**
The commutator consists of a commercial RF slip-ring with a top (static) and a bottom (rotating) SMA connector. The bottom connector is actively rotated by a stepper motor via a pair of custom 3D-printed gears, one of which is attached to the slip ring, and one to the motor axle. A custom driver board interfaces with the host PC with a standard USB interface and generates appropriate motor control signals as instructed by the host PC. Capacitive touch buttons on the outer side of the printed circuit board allow for manual operation of the commutator. The entire system is powered by the 5V provided from the USB interface.

**Electrode Interface Boards**
To support the use of the headstages, we have created electrode interface boards (EIB) adapting for interfacing with tetrodes drives[44], or other electrode or drive implants that use omnetics connectors. The designs primarily cater to tetrode use, but the schematics can be easily adapted to different PCB form factors for other probe designs. In addition to recording electrodes, all EIBs include outputs from the headstage for onboard LED drivers and electrical microstimulators which can be connected to drivable LED pigtails or electrodes, respectively. Electrodes can be soldered to through-holes or compression-connected using conical, gold-plated pins (available from NeuraLynx). These boards contain four copper layers. Sensitive analog traces are routed in internal layers. Top and bottom layers are connected to system ground providing near complete Faraday shielding of analog traces.

### Mouse behavior and neural recordings

All mice used for chronic electrophysiology verification were used for a separate study[61] before use in the experiments described here. Non-implanted mice used for behavior verification were handled under the same protocol as outlined here, but with no surgical intervention.
**Drive implants:** Light weight drive implants with 16 movable tetrodes were built as described previously[44,61]. The tetrodes were arranged in an elongated array of approximately 1250x750 µm, with an average distance between electrodes was 250 µm. Tetrodes were constructed from 12.7 µm nichrome wire (Sandvik – Kanthal, QH PAC polyimide coated) with an automated tetrode twisting machine[68] and gold-electroplated to an impedance of approximately 300 KΩ.

**Surgery:** Mice (C57BL/6 RRID: IMSR_JAX:000664) were aged 8-15 weeks at the time of surgery. Animals were housed in pairs or triples when possible and maintained on a 12-h cycle. All experiments were conducted in accordance with the National Institutes of Health guidelines and with the approval of the Committee on Animal Care at the Massachusetts Institute of Technology (MIT). All surgeries were performed under aseptic conditions under stereotaxic guidance. Mice were anesthetized with isofluorane (2% induction, 0.75–1.25% maintenance in 1 l/min oxygen) and secured in a stereotaxic apparatus. A heating pad was used to maintain body temperature, additional heating was provided until fully recovered. The scalp was shaved, wiped with hair-removal cream and cleaned with iodine solution and alcohol. After intraperitoneal (IP) injection of dexamethasone (4 mg/kg), Carprofen (5mg/kg), subcutaneous injection of slow-release Buprenorphine (0.5 mg/kg), and local application of Lidocaine, the skull was exposed. The skull was cleaned with ethanol, and a thin base of adhesive cement (C&B Metabond and Ivoclar Vivadent Tetric EvoFlow) was applied. A stainless-steel screw was implanted superficially anterior of bregma to serve as electrical ground. A 3 mm craniotomy was drilled over central midline cortex, a durotomy was performed on one side of the central sinus, and tetrode drives[44] were implanted above Retrosplenial cortex, at around AP -1.25 to -2.5 mm and ML 0.5 mm, with the long axis of the tetrode array oriented AP, and the tetrode array tilted inwards at an angle of ~15-20°, and fixed with dental cement. The ground connection on the drive was connected to the ground screw, and the skin around the drive implant was brought over the base layer of adhesive as much as possible to minimize the resulting wound margin, sutured, and secured with surgical adhesive.

At the time of implant surgery, only two of the tetrodes were extended from the drive to serve as guides during the procedure. All other tetrodes were lowered into superficial layers of cortex within 3 days post-surgery. Mice were given 1 week to recover before the start of recordings.

**Chronic Electrophysiology:** After implant surgery, individual tetrodes were lowered over the course of multiple days until a depth corresponding to cortical layer 5 was reached and spiking activity was evident. Data were acquired with an Open Ephys[69] ONIX prototype system at 30 kHz using the Bonsai software[54]. The tether connecting the mouse headstage to the acquisition system was routed through a motorized commutator above the arena and was counterbalanced via a segment of flexible rubber tread.

**Spike sorting:** Voltage data from the 16 tetrodes, sampled at 30 kHz were band-pass filtered at 300-6000 Hz, and a median of the voltage across all channels that were well connected to tetrode contacts was subtracted from each channel to reduce common-mode noise such as licking artifacts.

Spike sorting was then performed per tetrode using Mountainsort[70] (https://github.com/flatironinstitute/mountainsort_examples), and neurons were included for further analysis if they had a noise overlap score below 0.05, an isolation score > 0.75 (provided by Mountainsort[70]), a clear refractory period (to ensure spikes originated from single neurons), and a spike waveform with one peak and a clear asymmetry (to exclude recordings from passing axon segments), and a smooth voltage waveform and histogram (to exclude occasional spike candidates driven by electrical noise). Units were not excluded based on firing rates, tuning, or any higher order firing properties.

**Miniscope recording:** Basic surgery procedures were the same as described for drive implant surgeries. Mice were then unilaterally microinjected with 500 nL of AAV1-syn-jGCaMP7f-WPRE12 (Addgene) at 50 nL/min using the stereotactic coordinates: −2.1mm posterior to bregma, 1.5mm lateral to midline and −1.5 mm ventral to skull surface. Two weeks later, a gradient refractive index lens (GRIN) was implanted above the previous injection site. A 1.5mm diameter circular craniotomy was centered at the previous virus injection site. Artificial cerebrospinal fluid (ACSF) was repeatedly applied to the exposed tissue to prevent drying. The cortex directly below the craniotomy was aspirated with a 27-gauge blunt syringe needle attached to a vacuum pump. The GRIN lens (1.0mm diameter, 0.5 pitch, 4.0mm length, Inscopix) was slowly lowered with a stereotaxic arm above CA1 to a depth of 1.45mm ventral to the surface of the skull. The GRIN lens is then fixed to the skull using cyanoacrylate glue and dental cement. Two weeks later, a small rectangular baseplate was cemented onto the animal's head atop the previously formed dental cement. During imaging, the microendoscope (UCLA miniscope V4) was fixed in place inside the baseplate. The microscope's focus was adjusted electronically before recording to ensure the cells were in focus.

**Neuropixels recording:** To validate the use of ONIX with Neuropixels probes, a separate experiment was carried out at the Allen Institute following protocols approved by the internal IACUC under an assurance with the NIH Office of Laboratory Animal Welfare. A ChAT-IRES-Cre transgenic mouse (Jackson Labs) was implanted with a titanium headframe, and most of the left parietal skull plate was removed and replaced with a 3D printed cap. After four weeks of recovery, the protective silicone elastomer was removed from the skull cap under isoflurane anesthesia and replaced with a 1 mm thick layer of Duragel (DOW DOWSIL™ 3-4680). In the same procedure, a silver ground wire was inserted through one of the anterior holes in the 3D printed skull cap until it was just touching the brain surface. Starting on the following day, the mouse was habituated to head fixation on the recording rig for three consecutive days. On the day of the recording, a Neuropixels 1.0 probe connected to an ONIX headstage was inserted through one of the holes in the skull cap at a rate of 200 µm/min to a depth of 2.5 mm. After waiting 5 minutes for the probe to settle, 384 channels of AP band and LFP band data were recorded for 15 minutes at 30 kHz using Bonsai. The recording was made in external reference mode, with the Neuropixels ground and reference soldered together. Prior to visualization, the raw AP band data was phase shifted, high-pass filtered (300 Hz cutoff), and median subtracted using SpikeInterface[71].

**Behavioral experiment hardware:** Behavior was carried out in a hexagonal arena of 1.6m diameter. Individual floor tiles varied in height in a randomly chosen pattern to give mice the ability to run and jump across gaps spontaneously. The floor tiles were made of Styrofoam, painted with a water-resistant acrylic primer. All behavioral experiments were conducted in Bonsai[54], the code for conducting the recordings is available on the ONIX GitHub repository (https://open-ephys.github.io/onix-docs/Software%20Guide/Bonsai.ONIX/index.html).

**Behavioral analysis:** For non-implanted animals and comparison between these animals and implanted animals, marker-less 3D tracking[18] from an array of 5 cameras (Suppl. Fig. 5) was used to measure head position and orientation in 3D (Fig.2). For implanted animals where no direct comparison to non-implanted mice was performed (Fig.3), head-position and 3D-tracking data from the headstage were aligned in time and re-sampled to 100 Hz for further analysis. In all cases, distributions for running speed, head posture, and location were plotted from the resulting 3D-position and pitch/yaw data. For comparisons between implanted and non-implanted animals, occupancy was computed in a 20x20 grid spanning the entire maze, excluding periods where the mouse was stationary, and occupancy distributions were compared excluding one home position where each mouse spent a higher proportion of time. Maze occupancy was compared by computing their Shannon entropy and re-sampling in time bins of ~1 minute to compute bootstrap samples. The entropy of the occupancies was statistically indistinguishable within 4 hours of standard tether vs. 4 hours of ONIX tether data, with median entropies of 0.273 vs. 0.287 bit (normalized against uniform distributions with equal N of bins, see Voigts et al 2022 [61] ) and 95% confidence bounds of [0.208,0.369] vs. [0.224,0.383] bit. The entropy of the spatial occupancy for the classic headstage epochs had a median of 4.217 bit with a confidence interval of [3.998, 4.381] bit. Heading distributions (Fig. 2c) were compared with the same method, but in 40x40 bins spanning 0-360° yaw and ±45°pitch.
